# Statistical learning quantifies transposable element-mediated cis-regulation

**DOI:** 10.1101/2022.09.23.509180

**Authors:** Cyril Pulver, Delphine Grün, Julien Duc, Shaoline Sheppard, Evarist Planet, Raphaël de Fondeville, Julien Pontis, Didier Trono

**Affiliations:** School of Life Sciences, Swis Federal Institute of Technology Lausanne (EPFL), 1015 Lausanne, Switzerland; Swiss Data Science Center, Swiss Federal Institute of Technology Lausanne (EPFL), 1015 Lausanne, Switzerland

**Keywords:** transposable elements, transcription factors, gene regulation, cis-regulatory elements, embryogenesis, gene regulatory networks, epigenomics, transcriptomics, regulatory motif activity, RNA-seq, CRISPRi

## Abstract

**Background:** Transposable elements (TEs) have colonized the genomes of most metazoans, and many TE-embedded sequences function as cis-regulatory elements (CREs) for genes involved in a wide range of biological processes from early embryogenesis to innate immune responses. Because of their repetitive nature, TEs have the potential to form CRE platforms enabling the coordinated and genome-wide regulation of protein-coding genes by only a handful of trans-acting transcription factors (TFs).

**Results:** Here, we directly test this hypothesis through mathematical modeling and demonstrate that differences in expression at protein-coding genes alone are sufficient to estimate the magnitude and significance of TE-contributed cis-regulatory activities, even in contexts where TE-derived transcription fails to do so. We leverage hundreds of overexpression experiments and estimate that, overall, gene expression is influenced by TE-embedded CREs situated within approximately 200kb of promoters. Focusing on the cis-regulatory potential of TEs within the gene regulatory network of human embryonic stem cells, we find that pluripotency-specific and evolutionarily young TE subfamilies can be reactivated by TFs involved in post-implantation embryogenesis. Finally, we show that TE subfamilies can be split into truly regulatorily active versus inactive fractions based on additional information such as matched epigenomic data, observing that TF binding may better predict TE cis-regulatory activity than differences in histone marks.

**Conclusion:** Our results suggest that TE-embedded CREs contribute to gene regulation during and beyond gastrulation. On a methodological level, we provide a statistical tool that infers TE-dependent cis-regulation from RNA-seq data alone, thus facilitating the study of TEs in the next-generation sequencing era.

## Introduction

The development and function of complex organisms relies on the tight regulation of protein concentrations at cellular and tissue levels. Protein levels partially depend on cellular mRNA concentrations, in turn controlled by dynamic rates of gene expression. Cis-regulatory elements (CREs) are non-coding sequences that modulate the transcription of nearby genes in response to signaling cues, thereby contributing to the control of gene expression. Functionally, CREs operate through transcription factor (TF) recruitment and local chromatin remodeling [1]. Importantly, sequence specific TF-DNA binding allows for the simultaneous regulation of arbitrarily distant genes flanked by CREs carrying analogous TF binding sites (TFBS). Conceptually, the functional interactions implicating CREs, their target genes and their TF controllers form graph-like representations of the gene expression machinery known as gene regulatory networks (GRNs) [2, 3]. Typically, one may represent CREs as edges connecting two types of nodes: TFs and the protein-coding genes they regulate. Cell-state and tissue-specific transcriptional programs - defined by specific sets of expressed TFs and accessible CREs - are thereby depicted by distinct GRN topologies. For example, the GRN of so-called “primed” human embryonic stem cells (hESCs), which resemble epiblast cells of the post-implantation embryo, is characterized by the expression of OCT4, NANOG and SOX2. These TFs cooperate and bind to specific CREs, thereby maintaining the gene expression program of pluripotency [4]. Changes in TF expression can polarize cells towards a different state and thus alter GRN topology. For example, induced expression of Krüppel-like factor family (KLF) members in primed hESCs alters their GRN towards one resembling that of preimplantation-like “naïve” hESCs notably characterized by increased chromatin accessibility [5, 6].

Whereas the repertoire of expressed TFs and accessible CREs vary across cell states within one organism, the genomic location of CREs with respect to their target genes varies across species. In fact, it has long been recognized that organisms evolve primarily through the emergence, spread and reorganization of CREs [2, 7] rather than through mutations affecting protein-coding genes. In particular, TFs remain highly conserved across species though exceptions to this tenet exist [8]. In other words, the genomic reorganization of CREs and therefore the emergence of species-specific GRN topologies underlie the evolution of species. Thus, understanding the mechanisms by which GRNs become modified throughout evolution is a central question in biology. GRNs may evolve through large-scale chromosomal or even genome-wide duplication events followed by divergence and specialization of the henceforth redundant regulatory sub-networks. However, large-scale duplications are too coarse to account for the fine-grained nuances in CRE compositions observed upon comparing the genomes of distinct species. Due to their important contribution to the size of most metazoan genomes, their intrinsic ability to recruit TFs and their potential for rapidly spreading ready-to-go regulatory modules throughout the genome of their host, transposable elements (TEs) have gained attention as a potential source of CREs [2, 9, 10].

TEs form a collection of genetic entities that autonomously or collectively code for the factors essential to their own mobility, a process known as transposition. Endogenous retroelements (EREs) propagate through retrotransposition, a copy- and-paste mechanism entailing the reverse transcription of an RNA intermediate encoded within the ERE sequence itself. In agreement with the replicative nature of retrotransposition, EREs constitute the vast majority of the approx. 4.5 million readily recognizable TE-derived sequences that contribute more than half of the human genome DNA content [11, 12]. In contrast, DNA transposons propagate through a non-replicative cut-and-paste process and rely on genome replication to accumulate copies [13]. Both EREs and DNA transposons are further segregated into superfamilies and subfamilies [14]. TE subfamilies are sets of integrants that generally share a high degree of sequence similarity and use the same mechanism for transposition [15]. Britten and Davidson demonstrated long before the Next Generation Sequencing era that most metazoan genomes were replete with repetitive sequences, some of which emerged in recent evolutionary times [2]. Hence, they reasoned that repetitive DNA may form a pool of potential CREs whose cycles of expansion followed by purifying selection fuels GRN evolution. Consistent with this model, binding sites of conserved TFs and open chromatin regions are enriched in evolutionarily young TE subfamilies, in particular in embryonic stem cells (ESCs), and more occasionally in cancer cell lines and lymphoblastoid tissues [16, 17, 18, 19, 20]. Moreover, multiple functional studies support the regulatory potential of TEs, including evolutionarily recent integrants. For example, the majority of genes deregulated in human but not in mouse embryonic stem cells (mESCs) upon knockdown of the master regulator of pluripotency OCT4 are associated with EREs of the ERV1 family, for which an enhancer activity was confirmed by reporter assay [21]. As well, the majority of species-specific enhancers in mouse and rat trophoblast stem cells overlap species-specific TE subfamilies, and a mouse specific subfamily (RLTR13D5) exhibits trophoblast stem cell-specific enhancer activity in a reporter assay [22]. Finally, the genetic excision of primate-specific MER41B integrants thwarts the functionality of a key innate immunity signaling cascade [23] and hundreds of genes including stemness maintainers are downregulated upon epigenetic repression of the hominoid-specific SVA and LTR5-Hs subfamilies in hESCs [5]. Together, these case studies suggest that evolutionarily recent EREs spread CREs upon which natural selection may act to fine-tune the GRNs of critical physiological processes such as embryogenesis and innate immunity [9, 12, 10]. Despite accumulating evidence that some TE subfamilies form sets of functional CREs, no well-defined and genome-wide statistical framework has been proposed to estimate whether and how much TEs influence the expression of protein-coding genes. In addition, the identification of TE-embedded CREs currently relies on the genomewide profiling of epigenomic hallmarks such as histone modifications, TF binding, enhancer RNA (eRNA) production and endonuclease or tagmentation sensitivity [16, 21, 24, 19, 20]. While these assays are instrumental to characterize exhaustively the involvement of TEs as CREs under specific biological contexts, performing them in pair with RNA-seq significantly increases experimental costs as well as the biological material required prior to sequencing. Thus, a statistical framework based on RNA-seq and capable of estimating which TE subfamilies serve as CREs would benefit the gene regulation research field for hypothesis generation and data interpretation at negligible additional costs.

The hypothesis that TEs influence the expression of protein-coding genes at the subfamily level has a corollary: one should be able estimate the contribution of TEs to the expression of protein-coding gene by formulating a TE-centric mathematical model of gene regulation from basic principles of gene regulation. Analogous models have been developed to estimate the regulatory activity of TFBS motifs using transcriptomic data [25, 26, 27, 28]. These statistical approaches assume that DNA motifs or sequences - typically corresponding to TFBS - may regulate all promoters within which they are present with a quantitatively similar effect on gene expression. By analogy, as TEs evolved to attract the TFs necessary to trigger their own mobility, they can be conceptualized as larger regulatory sequences denoted as TE-embedded regulatory sequences (TEeRS). Thus, we took inspiration from the model of gene regulation championed by Britten and Davidson [2] and hypothesized that phylogenetically related TE integrants may attract similar sets of transcriptional regulators and hence bear a similar regulatory influence on protein-coding genes located in their vicinity. Our system, coined *craTEs* (cis-regulatory activities of Transposable Element subfamilies), models variations in gene expression as a linear function of the susceptibility of protein-coding genes to the cis-regulatory activity of TE subfamilies. Here, we define activity as the variation in gene expression which can be attributed to the presence of integrants belonging to a set of phylogenetically related TEs within cis-regulatory distance of the gene promoter. In this study, we assume a priori that TE subfamilies form said sets. *craTEs* thereby enables the identification of cis-regulatory TE subfamilies from RNA-seq data alone and does so in a manner dependent on the expression profile of protein-coding genes. Thus, *craTEs* adheres to a strict definition of cis-regulatory activity which requires an associated change in gene expression, in contrast with approaches relying solely on the profiling of biochemical activity at TE loci [17, 18, 24, 19, 20].

In this study, we start by showing that *craTEs* accurately identifies cis-regulatory TE subfamilies from RNA-seq data alone. We demonstrate that it achieves this feat agnostically with respect to TE-derived transcription, with increased statistical power compared with standard enrichment-based approaches, and in cases where changes in transcription at the corresponding subfamilies remain undetectable. We then leverage *craTEs* in conjunction with a large-scale TF perturbation RNA-seq dataset to estimate the maximal genomic distance up to which cis-regulatory Tes significantly contribute to the regulation of transcription genome-wide. Using the same dataset, we then identify novel regulatory links between TF expression and cis-regulatory TE activities. Finally, we verify that *craTEs* detects biologically relevant regulatory phenomena by performing DNA binding and histone mark profiling experiments. Overall, we present and validate *craTEs*, a simple mathematical model of TE-dependent gene regulation. *craTEs* recapitulates the findings of landmark case studies of TE-dependent cis-regulation and suggests previously unappreciated regulatory ties implicating TFs and TEs. These results support a model of GRN evolution whereby the spread of TEs provides an important supply of raw regulatory materials.

## Results

### *craTEs* models variations in gene expression as a linear combination of TE-encoded cis-regulatory elements

Using RNA-Seq data, we aimed to systematically uncover TE subfamilies that regulate the expression of protein-coding genes in cis. Integrants of the same TE sub-family share a high level of sequence similarity. Thus, they are predicted to exert a similar cis-regulatory influence on protein-coding genes located in their vicinity. We assumed that the subfamily composition of TE integrants located within cis-regulatory distance of protein-coding genes contributes to a significant fraction of the variation in gene expression (fig.1A) [26, 28]. As a first approach, we set this distance to 50kb since differentially expressed (DE) genes were found to be enriched within this range of epigenetically perturbed cis-regulatory LTR5-Hs and SVA TE subfamilies [5].

Considering two experimental conditions denoted as 1 and 2, for example “control cells” and “cells with transgene overexpression”, we modeled the variation in gene expression Δ*E*^′^ of each of the *p* protein-coding gene as a linear combination of the per-subfamily TE integrant counts *N*_*pm*_ located within cis-regulatory distance of its promoters. *N*_*pm*_ represents the regulatory susceptibility [26] of gene *p* to TE sub-family *m*. We trade biological complexity for statistical simplicity by treating me bers of the same TE subfamily as “regulatory black boxes” of equal cis-regulatory potential. A well-known caveat of currently available ERE annotations is that integrants are often fragmented into multiple sequences [12, 29], causing an artificial inflation of *N*_*pm*_ and potentially deteriorating model performances. We therefore merged closely located ERE fragments of the same subfamily into single ERE integrants. LINEs, LTRs and SVAs were particularly prone to spurious fragmentation (fig. 1B), with numbers of integrants dropping by 8.3%, 7,9% and 15% respectively after merging. To define the regulatory susceptibility *N*_*pm*_ of each gene *p* to each TE subfamily *m*, we counted the number of integrants of subfamily *m* falling within cis-regulatory distance of the promoters of *p*. We found that between 45.9% (LTRs) and 72.5% (SVAs) of all integrants were located within cis-regulatory distance, i.e. 50kb up/downstream, of at least one protein coding gene promoter (fig. 1B). In rare instances, TEs overlap gene exons. Since these are used to quantify RNA-seq reads, this may introduce a spurious association between the presence of an annotated TE integrant and gene expression. We addressed this by excluding TEs overlapping exons from the set of putatively cis-regulatory TEs susceptible to regulate the corresponding gene. Finally, we chose to emphasize TE-driven cis-regulation dependent on distal sequences, i.e. located more than 1.5kB up/downstream of a transcription start site, as the role of TEs as alternative promoters has been extensively studied elsewhere [30, 31]. Thus, we prevented TEs overlapping with promoters of gene *p* from contributing to the set of regulatory susceptibilities *N*_*pm*_ (fig. 1A,B). The combination of the last two filtering steps excluded 1.2% of TEs found within cis-regulatory distance of protein-coding genes from *N* (fig. 1B).

**Figure 1.**
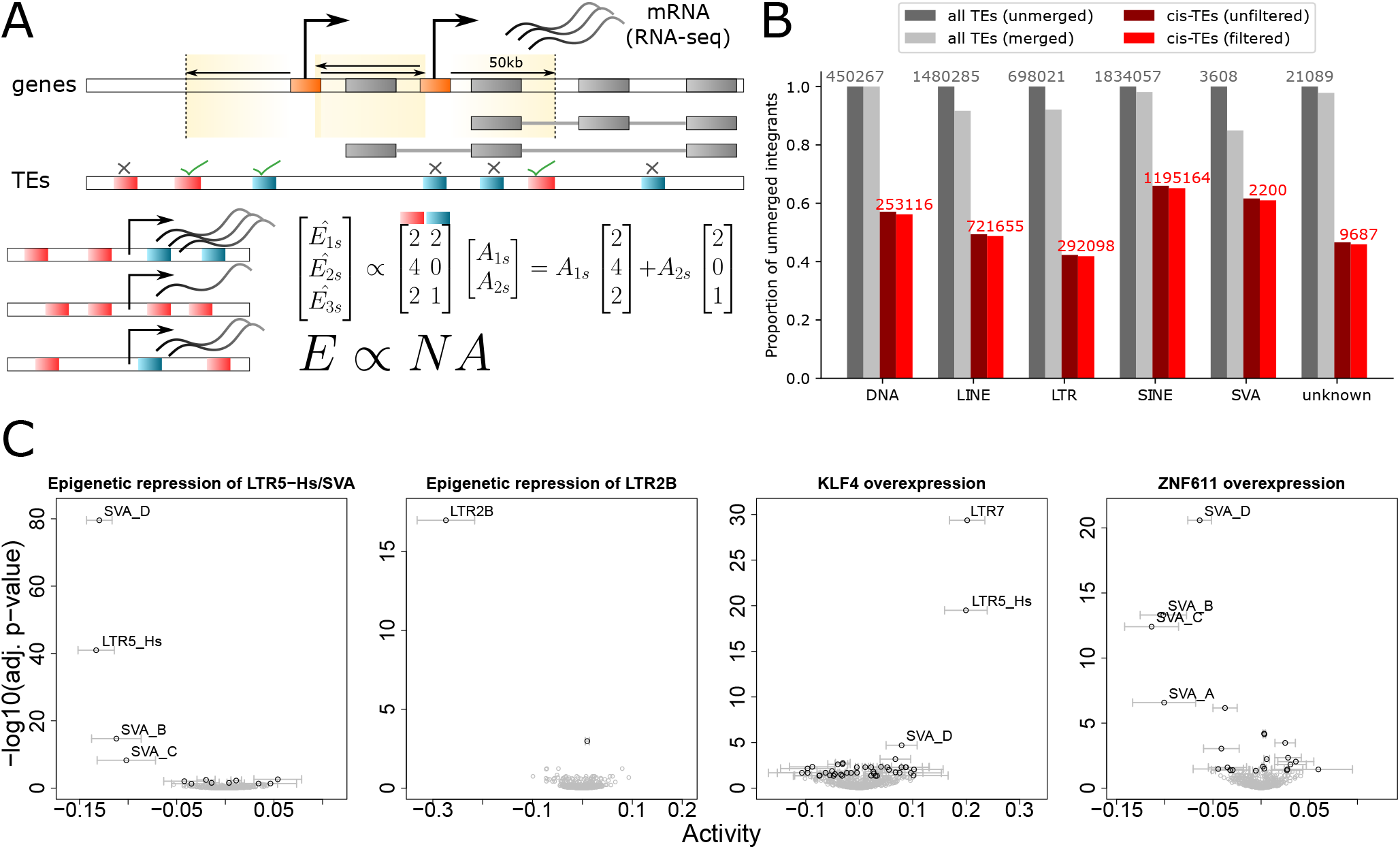
craTEs uncovers cis-regulatory TE subfamilies from RNA-seq. **A** Overview of the *craTEs* model. Differences in expression [log(TPM)] for protein-coding genes between treatment and control samples (columns of matrix E) are modeled as a linear combination of the per-subfamily TE counts found in the cis-regulatory region of each gene (columns of N). The cis-regulatory activities for each treatment vs control experiment (columns of A) are estimated by least squares. The cis-regulatory regions of each gene are defined as 50-kb long stretches of DNA 5’ and 3’ from promoter regions. Cis-regulatory regions exclude the exons and promoters of the genes they are assigned to. **B** Proportion of integrants remaining at each step of the construction of N with respect to the original number of TEs present in the annotation (indicated in grey). “All TEs” refers to all integrants found in the the TE database “Repeatmasker RELEASE 20170127” (number of unmerged TEs are indicated in grey). “cis-TEs” refers to integrants found in cis-regulatory regions before (“unfiltered”) and after (“filtered”, numbers indicated in red) removing those overlapping exons and promoters of the corresponding gene. **C** Four case studies exemplifying the estimation of the cis-regulatory activities of TE subfamilies from RNA-seq data. Black dots are TE subfamilies with significant (BH-adj. p-value ¡ 0.05) differences in activities between the treatment and control groups. 95% confidence intervals for the estimated cis-regulatory activities are shown as grey bars. Grey dots are TE subfamilies with non-significant differences in activities. From left to right: CRISPRi-mediated repression of LTR5-Hs and SVA integrants in naïve hESCs, *n* = 4 (4 treatment samples vs 4 control samples); CRISPRi-mediated repression of LTR2B integrants in K562, *n* = 2; overexpression of the pluripotency TF KLF4 in primed hESCs, *n* = 4; overexpression of the SVA-targeting KZFP ZNF611 in naïve hESCs, *n* = 2.

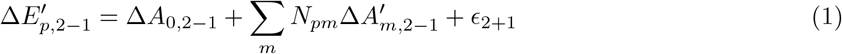

The main purpose of *craTEs* is the estimation of 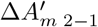 which we define as the difference in cis-regulatory activity exerted by subfamily *m* between conditions 1 and 2 (see equation 1). For the purpose of this study, we choose the convention that a positive cis-regulatory activity refers to an “enhancer-like” effect in condition 2 with respect to condition 1. Conversely, a negative cis-regulatory activity may reflect either the gain of a “silencer-like” effect or the loss of an “enhancer-like” effect in condition 2 versus condition 1. The cis-regulatory activity 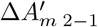 has an intuitive interpretation: it is the quantity in expression that would be gained by any gene in condition 2 with respect to condition 1 upon insertion of an integrant of subfamily *m* within cis-regulatory distance of one of its promoters. An independently and identically distributed Gaussian noise term centered around zero *ϵ*_2+1_ captures the variation in gene expression that is not accounted for by the linear model. *craTEs* estimates the vector of cis-regulatory TE subfamily activities 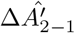 by minimizing the squared difference between the observed logged expression values 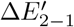 and those modeled as linear combinations of the columns of the susceptibility matrix *N*, containing the regulatory susceptibilities *N*_*pm*_. Further, *craTEs* assesses whether there is statistical evidence that 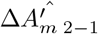 differs from zero: each component of 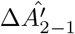 is tested against the null hypothesis 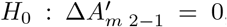, i.e that there is no difference in activity between condition 1 and 2 for that specific subfamily. This provides a measure of statistical significance for the estimated differences in TE-dependent cis-regulatory activities between conditions 1 and 2.

### *craTEs* uncovers cis-regulatory TE subfamilies from RNA-seq data

We then assessed the ability of *craTEs* to detect cis-regulatory TE subfamilies under controlled experimental settings. In particular, we leveraged two RNA-seq datasets derived from experiments in which specific TE subfamilies were epigenetically silenced, thus ablating their cis-regulatory effect on neighboring protein-coding genes [5, 32]. These datasets provide a biological “ground truth” against which we evaluated the output of *craTEs*. The targeted epigenetic repression of specific genomic loci was achieved by means of the CRISPR interference (CRISPRi) system [33]. CRISPRi relies upon a catalytically dead Cas9 domain (dCas9) that binds to DNA sequences complementary to user-defined guide RNAs (gRNAs). Once bound to the DNA, the dCas9-fused KRAB domain elicits the local deposition of repressive histone marks, thereby suppressing any enhancer activity exerted by the target site. As TEs of the same subfamily share a high level of sequence similarity, hundreds of related integrants can be targeted for silencing by only a handful of carefully designed gRNAs. We have previously shown that the hominoid-specific LTR5-Hs and SVA TE subfamilies serve as enhancers in naïve hESCs, and that this cis-regulatory activity can be ablated by CRISPRi [5]. We reanalyzed RNA-seq data from naïve hESCs where large fractions of the LTR5-Hs and SVA subfamilies were epigenetically silenced via CRISPRi. We applied *craTEs* to the vector 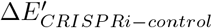 containing the differences in gene expression between the CRISPRi and control naïve hESCs. After correcting for multiple testing using the Benjamini-Hochberg procedure [34], we uncovered six TE subfamilies with significant differences in cis-regulatory activity (fig. 1C). Among these, LTR5-Hs, SVA B, C and D subfamilies displayed the largest and most significant absolute estimated cis-regulatory activities. The negative activity values reflect the abrogation of the enhancer effect exerted by LTR5-Hs and SVAs in naïve hESCs by the CRISPRi system. Thus, *craTEs* correctly inferred the loss of enhancer effect at the subfamilies targeted by CRISPRi and did so from the expression of protein-coding genes alone.

As it is well established that TEs are particularly active in hESCs [35, 6], we wondered whether *craTEs* would be able to recover TE-dependent cis-regulatory changes in other cellular contexts. A subset of LTR2B elements are marked by the enhancer histone mark H2K27ac in various leukemia cell lines, including in chronic myelogenous leukemia-derived K562 cells [32]. We used *craTEs* to estimate the differences in TE-driven cis-regulatory activities between K562 cells where LTR2B were repressed via CRISPRi and their control counterparts. *craTEs* correctly identified LTR2B as significantly less active in LTR2B-CRISPRi K562 cells compared to control K562 cells (fig. 1C). Thus, *craTEs* recovers TE-dependent cis-regulatory mechanisms beyond the context of hESCs.

Next, we empirically verified whether the ability of *craTEs* to detect changes in regulatory TE activity generalized beyond experiments of targeted TE repression via CRISPRi. TEs often exert cis-regulatory effects by serving as docking platforms for TFs. For example, the core pluripotency TF KLF4 is highly expressed in naïve hESCs, where it binds to LTR7, LTR5-Hs and SVA integrants [36, 5]. Interestingly, these subfamilies also display elevated levels of the enhancer histone mark H3K27ac in naïve hESCs. In contrast, primed hESCs generally express lower levels of KLF4 and TEs than their naïve hESCs [6, 5]. Using *craTEs*, we assessed the impact of KLF4 overexpression on TE-dependent cis-regulation in primed hESCs. *craTEs* identified LTR7, LTR5-Hs and SVA D as the most significant and highly activated TE subfamilies upon KLF4 overexpression, thereby recapitulating our previous findings [5] agnostically with respect to epigenomics data and TE-derived transcripts (fig. 1C). Interestingly, we previously observed that the KLF4-dependent enhancer activity of SVAs in primed hESCs did not correlate with increased SVA transcription but instead with an accumulation of H3K27ac enhancer histone marks at SVA integrants [5]. This suggests that *craTEs* detects TE-dependent cis-regulatory effects that would not be inferred from studying the variation in expression of TE integrants. Furthermore, overexpression of the repressive SVA-binder KRAB-zinc finger protein ZNF611 [8] in naïve hESCs abrogates the enhancer activity of SVAs [5]. We used *craTEs* to estimate the differences in TE-dependent cis-regulation between ZNF611-overexpressing naïve hESCs and control naïve hESCs. As expected, *craTEs* identified SVAs as the TE subfamilies with the most significant and largest absolute differences in cis-regulatory activity between control and ZNF611-overexpressing naïve hESCs (fig. 1C), with negative activity values reflecting the loss of enhancer effect at SVAs upon ZNF611 overexpression. Together, these results show that *craTEs* correctly identifies TE-dependent cis-regulatory activity changes beyond the context of targeted TE epigenetic perturbations and demonstrate its utility for identifying TE-dependent regulatory mechanisms under biological perturbations that affect TEs indirectly. In addition, *craTEs* identifies cis-regulatory TE subfamilies without resorting to mapping RNA-seq reads emanating from transcriptionally active TEs or performing epigenomics assays.

### *craTEs* outperforms enrichment approaches based on differential expression analyses

The notion that differences in gene expression may reveal candidate cis-regulatory TEs has already been exploited in previous studies [37, 5] though the statistical methodologies differ from *craTEs* in key aspects. More specifically, these methods identify cis-regulatory TEs through a two-step process. First, differentially expressed (DE) genes are identified through ad-hoc statistical methods [38, 39]. Then, per-subfamily scores for the enrichment of differentially expressed genes in the vicinity of TE integrants are computed. A high enrichment is reflected by a small probability (p-value) of finding more DE genes in the vicinity of a specific subfamily than the observed number of DE genes. We empirically compared the output of *craTEs* with that of the enrichment approach the RNA-seq dataset whereby LTR5-Hs/SVA were silenced via CRISPRi [5]. Using the enrichment approach, we found that DE genes whose expression fell under LTR5-Hs/SVA epigenetic repression (p-val ¡ 0.05, Fisher’s exact test) were significantly enriched in the vicinity of LTR5-Hs, SVA B and SVA D integrants (fig. 1B) (BH-adj. p-val ¡ 0.05, hypergeometric test). Note that the DE enrichment approach failed to detect the regulatory link between gene downregulation and the TE subfamily SVA C (adj. p-val = 1, hypergeometric test), whereas these were identified by *craTEs*. Moreover, when correcting for multiple testing during differential expression analysis (BH-adj. p-val ¡ 0.05, Fisher’s exact test), DE genes were enriched near SVA D integrants, but not LTR5-Hs or other SVA subfamilies (fig. 1C). Overall, this suggests that by pooling information across all genes, and not just DE genes, *craTEs* offers increased statistical power over classical DE enrichment approaches.

*craTEs* estimates cis-regulatory TE activities by considering expression variations across hundreds of protein-coding genes. Consequently, *craTEs* does not require replicates to estimate TE subfamily cis-regulatory activities. To illustrate this, we reanalyzed the RNA-seq data derived from the LTR5-Hs/SVA CRISPRi experiment [5]. We treated each pair of LTR5-HS/SVA CRISPRi and control samples as a single experiment, in effect ignoring the information provided by the replicate structure. We applied *craTEs* to each of the four replicates individually (fig. S1A). For LTR5-HS and SVA subfamilies, the estimated cis-regulatory activities are not surprisingly dropping and remain significant in all replicates, to the exception of SVA B and SVA C subfamilies for which only 1 out of 4, resp. 2 out of 4 replicates yield significant activities. Thus, whereas the discovery power of *craTEs* grows together with the number of replicates, the method can still uncover statistically significant changes in the cis-regulatory activities of TEs even in the absence of replicates. In contrast, any DE enrichment approach requires at least three samples due to the prerequisites of the DE analysis methods [38, 39]. In addition, *craTEs* not only quantifies the statistical significance of TE subfamily cis-regulatory activities but also provides a measure of the effect size through the estimated coefficient 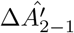 which can be interpreted as the difference in gene expression that would affect any gene upon insertion of a copy from the relevant subfamily within cis-regulatory distance. Overall, this case study suggests that *craTEs* is more powerful and more informative than DE-based enrichment approaches to discover cis-regulatory TE subfamilies. In addition, the superior statistical power of *craTEs* compared with DE gene enrichment approaches reinforces the notion that TEs act as cis-regulatory fine tuners, the dynamics of which may be overlooked when restricting the analysis to DE genes only.

### Influential TE-embedded cis-regulatory information resides up to 200kb from gene promoters

In a first implementation of *craTEs*, we defined cis-regulatory regions as 50kb-long stretches of DNA directly adjacent to the 5’ and 3’ sides of protein-coding gene promoters. Though informed by previous work[5], this choice of genomic distance was based upon data corresponding to LTR5-Hs and SVA subfamilies in hESCs only and may not reflect the general range of action of cis-regulatory TEs across all subfamilies and cellular contexts. We therefore modified the *craTEs* model to weight the regulatory influence of each integrant *i* on each gene *p* as a continuous and decreasing function of its distance *d*_*p,i*_ to the closest promoter of *p* (fig. 2A). We defined the regulatory susceptibility *N*_*pm*_ as:

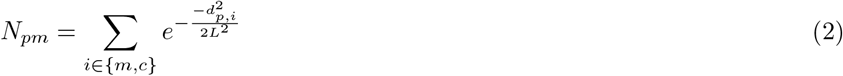

where each weight was computed using a gaussian kernel applied to the integrant-promoter distances *d*_*p,i*_. We considered all combinations of genes and integrants located on the same chromosome. Note that integrants falling within exons or promoters of *p* were excluded from *N*_*pm*_. We computed 11 susceptibility matrices *N* by varying the bandwidth of the gaussian kernel *L* between 1 kilobase (kb) and 10 Gigabases (Gb) thus spanning the entire range of possible cis-regulatory distances. Setting *L* to 1kb restricts cis-regulatory regions to the direct vicinity of gene promoters. In contrast, at 10 Gb, *L* exceeds the length of human chromosomes by two orders of magnitude, thus yielding nearly equal regulatory susceptibility scores across genes located on the same chromosome. We then tested which of these 11 matrices led to the smallest prediction error using 5-fold cross-validation. For the LTR5-Hs/SVA epigenetic repression, ZNF611 overexpression, KLF4 overexpression [5] and LTR2B epigenetic repression experiments [32], the validation error was minimized for *L* = 100kb or *L* = 500kb (fig. 2B). As 95% of the area under a gaussian curve is contained within two standard deviations from its mean, this suggests that TEs encode significant cis-regulatory information up to distances of approximately 200kb to 1 Million bases (Mb) from gene promoters. We note that errors estimated for small (1kb) and very large (*>*= 100Mb) values of *L* were unstable due to the high degree of collinearity between predictors. Indeed, a small *L* results in high numbers of zero-inflated columns in the *N* matrix. Conversely, very large values of *L* yield nearly equal weights for TE-gene pairs located on the same chromosome (fig.S2A). Both cases make the least squares problem ill-posed by making the matrix *N* singular.

**Figure 2.**
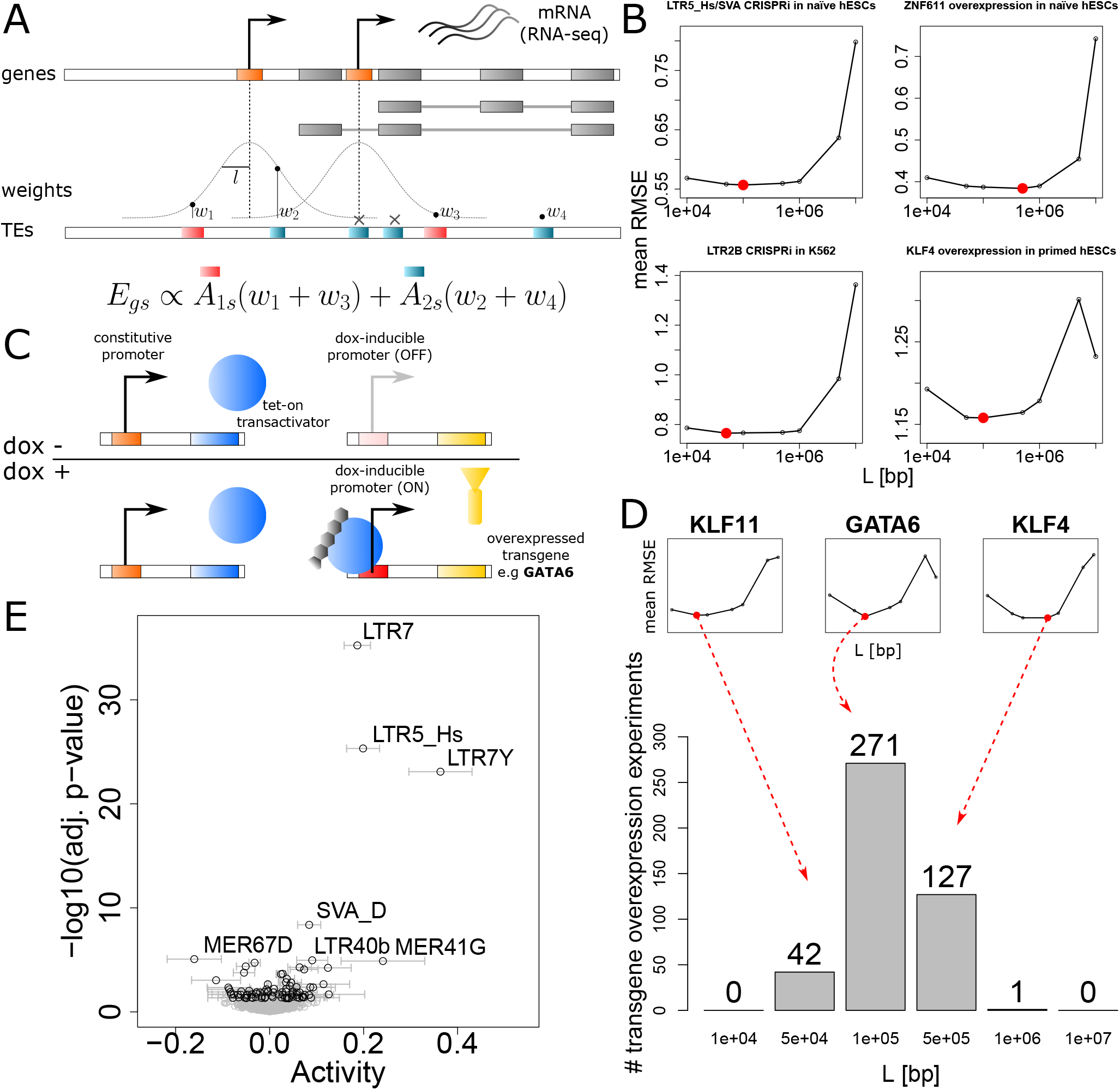
Influential TE-embedded cis-regulatory information resides up to 200kb from gene promoters. **A** Overview of the weighting process whereby the cis-regulatory influence of TEs decreases as a function of the distance to the closest promoter. The scheme depicts a protein-coding gene with two alternative promoters (in orange), coding for two alternative isoforms (in grey). Gaussian kernels with a maximum value of 1 and of varying bandwidth *L* are centered on each promoter. Before being added to the corresponding element in the matrix *N*, each TE is weighted as a function of its distance to the closest gaussian kernel. TEs overlapping exons and promoters are still excluded. **B** Optimizing the kernel bandwidth *L* for each of the experiments described in Figure 1C. The root mean-squared error (RMSE) was computed for each validation fold and averaged across the five folds. **C** Overview of the experimental design of the hESC “perturbome” [40]. Stable hESC cell lines carrying a stably integrated dox-inducible transgene overexpression construct were established from individual cells. In each of the 441 transgene overexpression experiments, dox-treated samples (dox+) are compared to the same cell line in the absence of dox (dox-). Note that the number of replicates per experiment varies. **D** Histogram depicting the number of times each gaussian kernel bandwidth *L* led to the smallest mean validation RMSE in a 5-fold cross-validation scheme for the 441 transgene overexpression experiments. Individual mean RMSE estimations for KLF11, GATA6 and KLF4 are showed as illustrative examples. **E** Estimation of the cis-regulatory activity of TE subfamilies upon KLF4 overexpression [5] using the matrix *N* computed with *L* = 100kb.

We wondered whether the optimal cis-regulatory bandwidths estimated from the four datasets treated thus far (fig. 2B) generalized to other TE subfamilies as well. We took advantage of a recently published RNA-seq dataset where hundreds of transgenes, mostly TFs, were overexpressed in primed hESCs through a dox-inducible system [40] (fig. 2C). We considered this dataset as a “perturbome” where each overexpressed transgene polarizes the primed hESC transcriptome towards a specific direction, e.g. towards the naïve hESC GRN or down a differentiation path. We used the same 5-fold cross-validation scheme to find the optimal value of *L* for each transgene overexpression experiment (fig. 2D). In 440/441 transgene overexpression experiments, the optimal bandwidth *L* took values between 50kb and 500kb. As most of the area below a gaussian curve is contained within two bandwidths from its mean, TE subfamilies encode cis-regulatory information up to distances comprised between 100kb and 1Mb from the promoters of protein-coding genes in hESCs. In 271/441 transgene overexpression experiments, *L* = 100kb led to the smallest cross-validation error, and thus can be chosen to weight cis-regulatory TE integrants such as to maximize predictability. As an illustration, with *L* = 100kb, a TE located 100kb away from a gene promoter receives a weight of 0.61 (fig. S2B). The weight drops down to 0.01 for a TE-promoter distance of 300kb for a virtually negligible contribution to the *craTEs* model.

A predictor matrix *N* based on TE contributions weighted by their distance to protein coding genes (fig. 2A) has two potential advantages over a predictor matrix *N* computed from hard distance thresholds as we did when first validating the discovery power of *craTEs* (fig. 1A). First, the quality of the predictors is likely to improve, as the optimal distance until which cis-regulation affects gene expression is estimated directly from the expression data. In other words, a continuous and decreasing weighting function may better represent the regulatory potential of TEs on protein-coding genes than a hard threshold approach. Second, as we require that each TE subfamily included in *N* sums up to a total regulatory potential greater than 150 (see the Methods section), the continuously decreasing weighting approach may allow for the inclusion of more TE subfamilies in the columns of *N*, leading to the discovery of previously overlooked significant cis-regulatory TE subfamilies. We used the KLF4 overexpression RNA-seq dataset we previously generated [5] to illustrate these points. We replaced the regulatory susceptibilities *N*_*pm*_ of matrix *N* computed according to a hard distance threshold (fig. 1A) with those corresponding to the same subfamilies, this time computed through TE-promoter distance weighting (fig. 2A, 2). Both models thus use the exact same number of predictors, i.e cover the same TE subfamilies. Running *craTEs* with the weighted matrix *N* computed with *L* = 100kb increased the fraction of gene expression variation explained from 4.6% to 5.6% compared to using the matrix *N* derived from a hard-thresholded cis-regulatory distance. As the number of predictors in *N* remained unchanged, this suggests that the distance weighting approach better approximates the cis-regulatory potential of TE subfamilies than the hard distance threshold approach. Next, we empirically evaluated whether allowing for the inclusion of TE subfamilies that passed the minimum per-subfamily regulatory potential with distance weighting (2) - but not with hard distance thresholding - would uncover additional biologically validated TE-dependent cis-regulatory changes. LTR7Y was identified as highly significantly activated upon KLF4 overexpression in hESCs (fig. 2E), in agreement with previously published results [5] while it was absent from the model depending on hard distance thresholding. In addition, though also absent from the hard distance thresholding model, the primate-specific MER41G subfamily was found as significantly and strongly activated in the distance-weighted model. To sum up, TE subfamilies typically encode cis-regulatory potential up to distances of approx. 200kb from the promoters of protein-coding genes, at least in the context of hESCs. This reinforces the notion that TEs form a layer of regulatory fine-tuners exerting a measurable impact on the expression of protein-coding genes.

### TFs controlling gastrulation and organogenesis promote the cis-regulatory activity of evolutionarily young TE subfamilies activated during pluripotency

Having validated the ability of *craTEs* to reveal bona fide TE-dependent cis-regulatory activities, we next aimed at characterizing the landscape of TF-induced TE-dependent cis-regulation in primed hESCs. As the epigenome of hESCs is markedly more open than that of differentiated cells [41], the number and strengths of the TF-TE regulatory interactions constituting the GRN of hESCs can be understood as upper bounds on those constituting the GRNs of differentiated tissues. We therefore applied *craTEs* to the “perturbome” dataset, where 441 transgenes, most of them TFs, were individually overexpressed in primed hESCs for 48 hours through a dox-inducible system (fig. 2C) [40]. Using the regulatory susceptibility matrix *N* computed according to the best performing cis-regulatory bandwidth (*L* = 100kb), we estimated the changes in cis-regulatory TE activities associated with each dox-induced transgene overexpression experiment (additional file 1). Dox-treatment alone and dox-induced GFP overexpression were not associated with any significant changes in cis-regulatory TE activities (fig. 3A) suggesting that neither the addition of doxycycline nor the metabolic cost entailed by strong transgene overexpression significantly altered the cis-regulatory activity of TE subfamilies. Furthermore, overexpression of the core pluripotency TF POU5F1 (also known as OCT4) was not associated with differential TE cis-regulatory activity (fig. 3A). This is consistent with the idea that overexpression of an already highly expressed gene such as POU5F1 in a cellular context that largely relies on it, i.e. primed hESCs, does not significantly alter TE-dependent cis-regulation. Together, these results suggest that TE-dependent cis-regulatory activities inferred from the remaining transgene overexpression experiments are not driven by technical factors inherent to the system used but induced by the overexpressed transgene itself.

**Figure 3.**
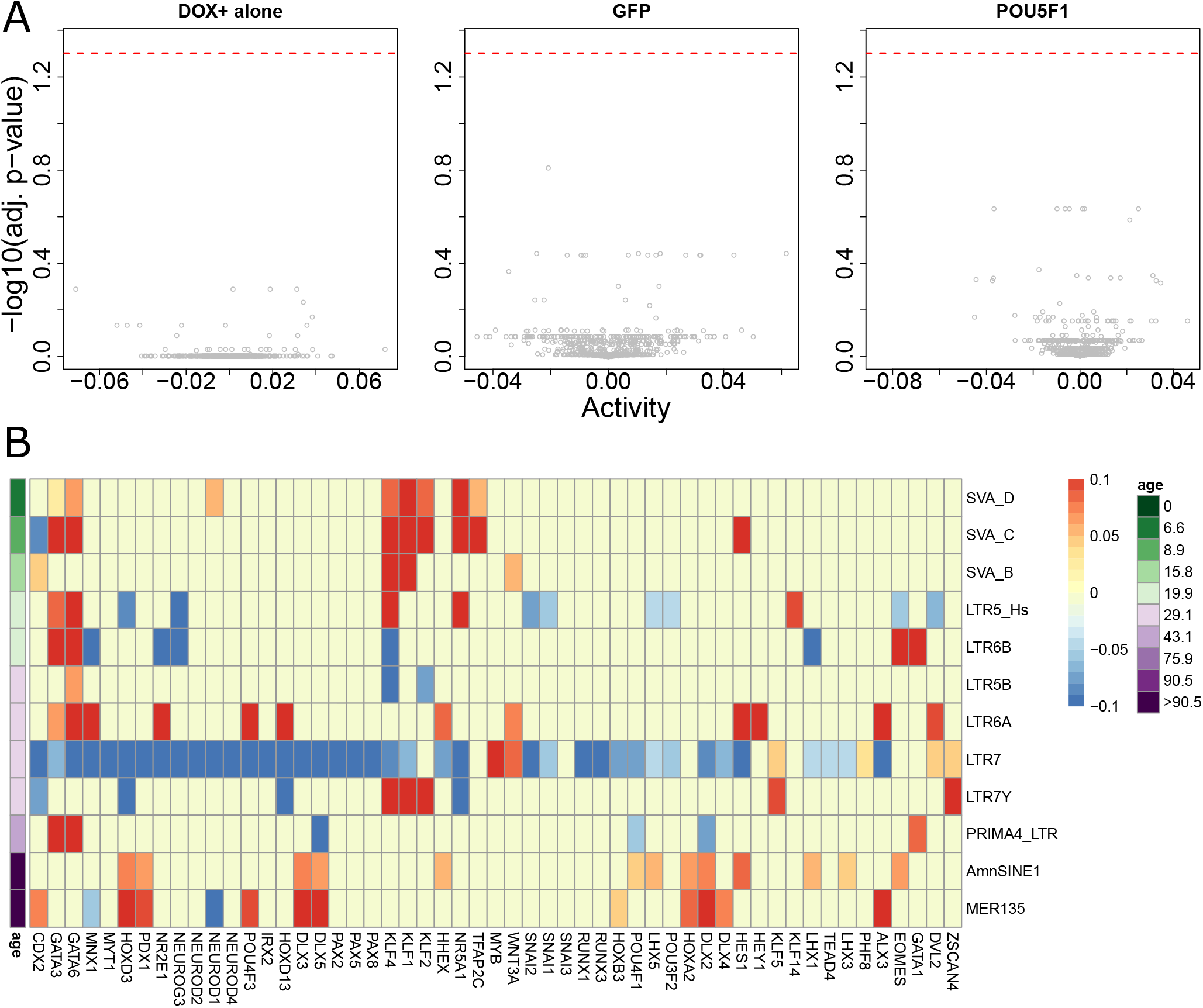
TFs controlling gastrulation and organogenesis promote the cis-regulatory activity of evolutionarily young TE subfamilies activated during pluripotency. **A** Estimation of the difference in TE subfamily cis-regulatory activities triggered by dox-treatment alone (left), dox-induced GFP overexpression (middle) and dox-induced POU5F1 overexpression (right) compared to the corresponding untreated cell lines. Dox-treatment was maintained for 48 hours. RNA-seq data obtained from a large scale transgene overexpression experiment in hESCs [40]. **B** TE subfamily cis-regulatory activities estimated from dox-induced TF overexpression experiments in primed hESCs. Cells were treated for 48 hours. The number of replicates for each condition varies. Non-significant differences in TE subfamily activities were set to 0, as the null hypothesis could not be rejected. The full heatmap was obtained by performing complete linkage hierarchical clustering based on the Euclidean distance computed between rows and columns of the matrix containing the statistical strengths of the differential activities (additional file 2). Selected TFs were ordered according to the dendrogram obtained from hierarchical clustering. TE subfamilies were ordered by evolutionary age in million of years, which was obtained from [5]. The color labeling of the estimated activities was saturated at |*A*| *<* 0.1.

To reveal how TE-dependent cis-regulation relies on TF overexpression in hESCs, we performed hierarchical clustering on the matrix containing the statistical strengths of the estimated cis-regulatory TE activities (additional file 2, fig, 3B). Overexpressed TFs of the same family tended to cluster together, e.g NEUROD1, NEUROD2, NEUROD3, NEUROD4; PAX2, PAX5, PAX8; SNAI1, SNAI2, SNAI3; RUNX1, RUNX3; HES1, HEY1; LHX1, LHX5, suggesting that commonalities in gene expression were partially mirrored by similar cis-regulatory TE activity patterns. Experiments where the core trophoblast TF CDX2 [42, 43] was overexpressed clustered away from all other experiments. This may reflect both the widespread rewiring of TE-dependent cis-regulation as primed hESCs differentiate towards trophectodermal cells [22] and a bias towards the detection of more differentially active cis-regulatory TE subfamilies in the CDX2 overexpression experiments due to a larger sample size compared to the other experiments.

The LTR7 subfamily clustered away from all other TE subfamilies (additional file 2) and was significantly less active in 209 out of 441 transgene overexpression experiments, making it the most frequently differentially active TE sub-family in this dataset. Among overexpressed transgenes tied to a decrease of LTR7-dependent cis-regulatory activity, we found multiple TFs involved in post-implantation developmental stages, e.g. the meso-endodermal master TF GATA6 [44] and several homeobox-domain-containing TFs including PDX1 and RUNX1. LTR7 cis-regulatory activity also decreased upon overexpression of organ and tissue-specific TFs, e.g NEUROD1, NEUROD2, NEUROD3, NEUROD4, MYT1, NR2E1, POU4F1, POU4F3, all involved in the formation of the nervous lineage [45, 46, 47, 48, 49]. Overexpression of TBX5, a key TF in the developing heart [50] also decreased the cis-regulatory activity of LTR7. Lastly, overexpression of TFs involved in the development and maintenance of the placenta e.g. CDX2, TEAD4 also led to a decrease of LTR7 cis-regulatory activity. Overall, inducing TFs tied to development and differentiation dampened the pluripotency-specific activity of LTR7 elements.

In contrast, we found rare transgenes (13/441) whose overexpression in primed hESCs led to an increase in LTR7 cis-regulatory activity (additional file 1, fig. 3B). Overexpressing KLFs, including KLF4, collectively increased the cis-regulatory activity of LTR7 and LTR7Y elements in agreement with previous studies characterizing KLFs as inducers of LTR7/LTR7Y enhancer activity in naïve hESCs [5]. Overexpression of MYB (also known as c-MYB), a TF involved in the maintenance of self-renewal in stem cells of the intestinal crypt, the bone marrow and the nervous system [52] as well as the formation of stem-like memory CD8 T cells [53], led to a marked increase in LTR7 cis-regulatory activity. Thus, MYB overexpression may reinforce self-renewal in the GRN of hESCs, a process tied to an increase in LTR7 cis-regulatory activity. More provocatively, this hints at the possible involvement of a MYB-LTR7 axis in the maintenance of self-renewal and stemness in adult cells and tissues. Our analysis thus suggests that a limited set of TFs linked to development and stemness may rely upon the enhancer potential of the LTR7 subfamily to establish, regulate and maintain these processes throughout development and adult life.

Other primate-specific TE subfamilies displayed partially overlapping patterns of cis-regulatory activity upon transgene overexpression. SVAs and LTR5-Hs were collectively activated by KLF4 and other KLFs (fig. 3B), consistent with previous work establishing the KLF4-dependent enhancer activity of these subfamilies in naïve hESCs [5]. Interestingly, the cis-regulatory activity of SVAs also increased upon overexpression of TFAP2C and NR5A1, both of which polarize hESCs towards the naïve state [54, 55]. Overall, these results suggest that recently emerged TE sub-families form functional collections of enhancer-like CREs during pre-gastrulation embryogenesis.

We then wondered whether the overexpression of transgenes necessary for embryonic development during and after gastrulation was associated with an increase in cis-regulatory activity in recently emerged TE subfamilies. Overexpression of the core meso/endodermal TF GATA6, as well as other GATA family members, increased the cis-regulatory activity of SVAs, LTR5-Hs and of LTR5B, thereby resulting in the activation of a TE-dependent cis-regulatory network partially reminiscing that of naïve hESCs [5]. This is surprising given that naïve hESCs resemble cells of the early blastocyst while GATA6 controls post-implantation developmental stages such as the formation of the mesoderm and the endoderm during gastrulation. Furthermore, GATA6 overexpression increased the cis-regulatory activity of additional primate-specific TE subfamilies including the ERV1 subfamilies LTR6A, LTR6B and PRIMA4-LTR. Of note, evidence for direct binding by multiple GATA family members derived from ChIP-seq was reported at LTR5-Hs, LTR6A, LTR6B and PRIMA4-LTR integrants [56], implying that the changes in cis-regulatory activity we observed may stem from direct binding of GATA TFs to these integrants. Lastly, overexpressing EOMES, a regulator of germ-layer formation and meso-endodermal differentiation [57], increased the cis-regulatory activity of LTR6B elements. Together, these results suggest that evolutionarily recent and pre-implantation specific TE subfamilies form sets of CREs that regulate the expression of protein-coding genes in cis well past the epiblast stage, including during and after gastrulation.

Older TE subfamilies that emerged prior to the speciation of primates also contribute to GRNs by donating CREs. Despite having spread before the speciation of amniotes hundreds of millions of years ago, AmnSINE1 elements are retained in the genomes of extant amniotes including humans and mice [58], and some AmnSINE1 elements were found to exert long-range enhancer effects on genes controlling brain development [59]. We observed that in primed hESCs, overexpression of several homeobox domain-containing TFs, e.g. RUNX1, a regulator of hematopoietic ontogeny [60], and PDX1, involved in pancreatic development [61], was associated with an increased AmnSINE1 cis-regulatory activity (fig. 3B). Interestingly, AmnSINE1 elements are enriched within active enhancers in epigenomes derived from fetal human cell lines [20]. Lastly, MER135, an ancient subfamily of currently unidentified origin [62] showed increased cis-regulatory activity upon overexpression of homeobox domain-containing TFs in primed hESCs (fig. 3B). More generally, these results hint that ancient TE subfamilies may retain their cis-regulatory potential at the subfamily level in extant species despite having colonized the genome of an evolutionarily distant common ancestor.

### Cis-regulatory activities are more pronounced at epigenetically active TEs

We showed that *craTEs* agnostically uncovers SVAs and LTR5-Hs as the subfamilies with the most significant and strongest loss of cis-regulatory activity upon CRISPRi-mediated epigenetic repression in naïve hESCs (Figure 1). However, it is highly likely that only a fraction of all integrants constituting a subfamily truly exert cis-regulatory effects. For example, integrants found within dynamic chromatin regions may be more differentially active than integrants located in stable chromatin regions. To empirically verify this hypothesis, we leveraged the matched epigenomic profiles matched to the LTR5-Hs/SVA CRISPRi RNA-seq dataset [5] and labeled the following integrants as “functional”: those overlapping with genomic coordinates where loss of chromatin accessibility (ATAC-seq) or gain of the repressive histone mark H3K9me3 (ChIP-seq) were detected upon epigenetic repression of SVAs and LTR5-Hs. Conversely and by complementarity, we considered all other integrants from these subfamilies as “non-functional”. We then expanded the weighted susceptibility matrix *N* (*L* = 100kb) column-wise by splitting TE sub-families into complementary functional and non-functional integrant subsets (fig. 4A). Finally, we used *craTEs* to jointly estimate the differences in cis-regulatory activity for functional and non-functional subsets of TE subfamilies upon epigenetic repression of LTR5-Hs and SVAs in naïve hESCs (fig. 4B). The functional subsets of SVAs and LTR5-Hs subfamilies displayed greater decreases in cis-regulatory activity upon epigenetic repression than complementary non-functional subsets. In addition, the estimated decreases in cis-regulatory activity were more pronounced for functional SVA and LTR5-Hs subsets than those estimated for the corresponding unsplit subfamilies.

**Figure 4.**
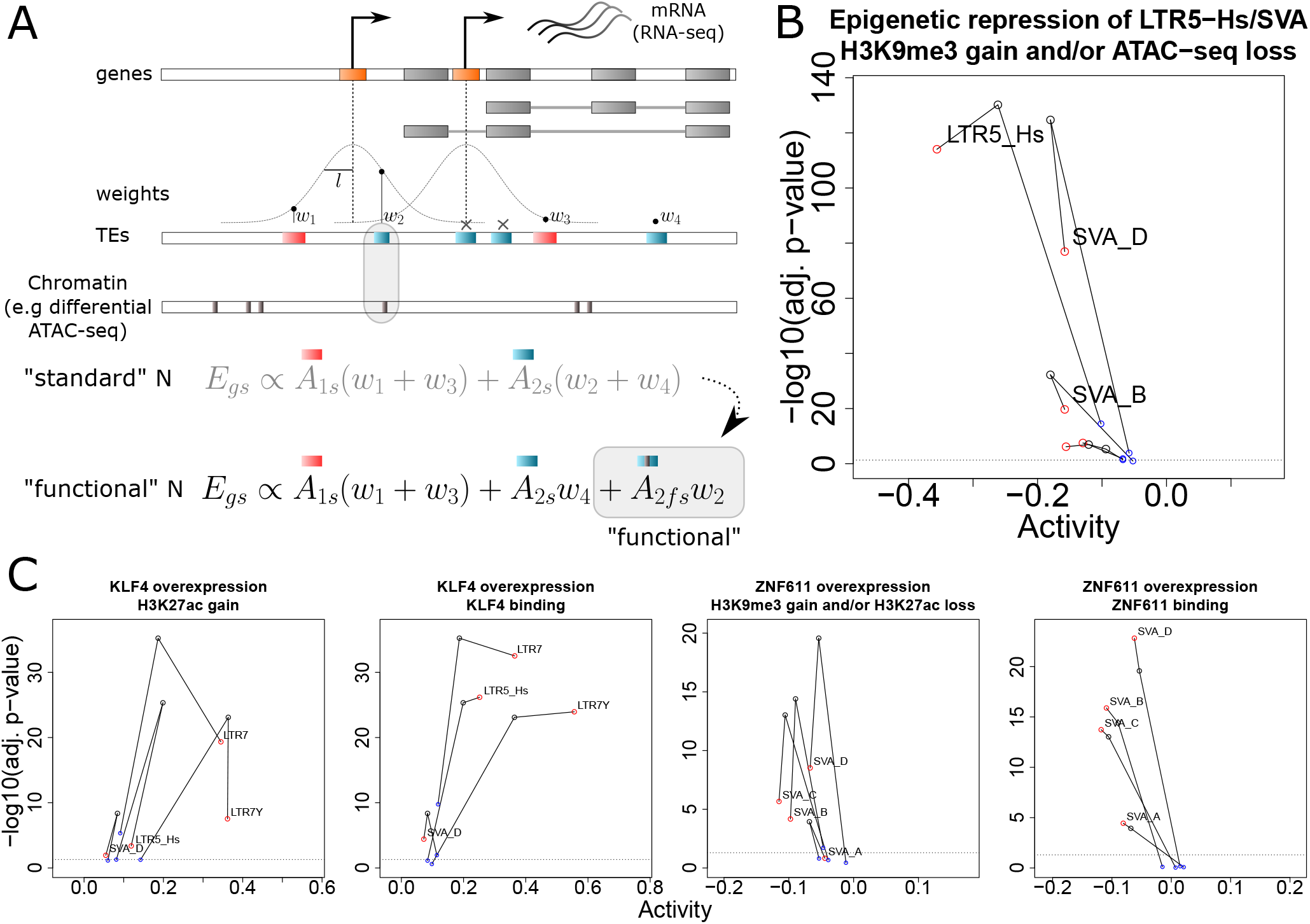
Cis-regulatory activities are more pronounced at epigenetically active TEs. **A** Overview of the procedure whereby TE subfamilies are split between so-called “functional” and “non-functional” fractions based on additional evidence, e.g differential chromatin accessibility. The regulatory susceptibility scores tying TE subfamilies to protein-coding genes are distributed between the functional and non-functional fractions of each TE subfamily, leading to an experiment-specific column-wise expansion of *N*. Concretely, functional and non-functional fractions of TE subfamilies are treated as independent TE subfamilies in the subsequent cis-regulatory activity estimation process. **B** Estimated cis-regulatory activities for the functional fraction (in red) and non-functional fraction (in blue) of LTR5-Hs and SVA subfamilies under CRISPRi-mediated epigenetic repression in naïve hESCs. The cis-regulatory activities for the unsplit subfamilies were estimated in a separated iteration of *craTEs*, using the standard *N* matrix, and are shown in black. The dotted line represents the significance threshold of BH-adjusted *p*.*val* = 0.05. Note that even though only selected subfamilies are plotted for clarity, all TE subfamilies were included in the fitting process. **C** Estimated cis-regulatory activities for the functional (in red) and non-functional fractions (in blue) of selected TE subfamilies according to definitions of the functional state that are either based on differential chromatin states (1st and 3rd panels from the left) or differential TF binding (2nd and 4th panels from the left) at integrants.

Finally, we leveraged the epigenomics-informed adaptation of *craTEs* to test whether differences in TF binding or histone marks could single-out integrants with detectable changes in cis-regulatory activity in the context of TF overexpression experiments. To this end, we completed the matched transcriptomics and histone profiles available for KLF4 and ZNF611 overexpression in hESCs [5] by generating ChIP-seq profiles against KLF4 and ZNF611. We used *craTEs* to estimate differences in cis-regulatory activity for functional integrant subsets defined according to differences in histone marks or TF binding (fig. 4C) and focused on the main cis-regulatory TE subfamilies identified under KLF4 and ZN611 overexpression in hESCs, namely LTR7, LTR5-Hs and SVAs. For both KLF4 and ZN611 overexpression experiments, histone mark-defined functional subsets had greater cis-regulatory activity than non-functional subsets, except for SVA-D under KLF4 overexpression. However, histone mark-defined functional subsets generally displayed only modest increases in cis-regulatory activity over unsplit subfamilies, at the cost of a marked decrease in statistical significance. In contrast, TF-bound integrants defined as functional displayed increased cis-regulatory activity with increased significance compared with the unsplit subfamily for all subfamilies except SVA-D under KLF4 overexpression. Thus, TF binding appears to better single-out bona fide cis-regulatory integrants than changes in histone marks. Together, these data suggest that TF binding may be better suited as a marker of cis-regulatory TE activity than changes in histone marks and empirically highlight that TE subfamilies can be further split into of active versus inactive sets of cis-regulatory integrants.

## Discussion

The notion that some TE-derived sequences behave as bona fide CREs is supported by an ever-growing number of reports, mostly relying on genome-wide profiles of promoter or enhancer specific histone marks. However, whether that biochemical activity should be interpreted as evidence for an evolutionary process fostering the emergence of collections of CREs to the benefit of the host, or instead as a byproduct of the so-called “selfish” tendency of TEs for genome invasion is subject to debate [9, 10, 14]. Still, if TEs truly spread functional CREs that become co-opted by the host through natural selection, one should at least be able to capture their effect on gene expression by modeling TE-dependent cis-regulation from basic gene regulation principles. Thus, we formulated *craTEs*, a model where differences in TE subfamily cis-regulatory activities are estimated in a single step from protein-coding gene expression. Using RNA-seq data derived from thoroughly characterized cases of TE-dependent cis-regulation, we showed that *craTEs* correctly identifies differentially active cis-regulatory TE subfamilies. Crucially, *craTEs* does not rely on TE-derived reads and is thus well-suited for the post-hoc analysis of standard RNA-seq count tables that did not take TE transcripts into account during feature quantification. In addition, *craTEs* was able to identify cis-regulatory TE subfamilies (SVAs) in RNA-seq datasets where no difference in transcriptional activity for these TE sub-families was previously detected [5]. These results suggest that TE subfamilies form at least partially consistent sets of CREs modulating gene expression in a coordinated fashion genome-wide and more generally that TEs spread highly resembling and functional cis-regulatory sites thereby supplying the raw materials critical to the evolution of coordinated gene regulation. We noticed that *craTEs* explains a fraction of the variation in gene expression ranging from approx. 3% to 10% in the experiments presented in fig. 1. This is comparable to the proportion of variance in gene expression explained by previously published linear models of gene regulation based on putative TF binding sites at core promoters [28]. In both cases, the low proportion of variance captured by the linear models is still sufficient for identifying significant and relevant regulatory mechanisms from transcriptomic data alone. However, while the true mathematical function underlying the regulatory mechanism at play is most likely non-linear [28], the linear model proposed in this work is still useful. Inferred activity coefficients are interpretable and well-established statistical tests exist to determine whether differences in cis-regulatory activities significantly deviate from zero. What is perhaps more impressive is that, in the context of this study at least, TE-dependent cis-regulation accounts for a fraction of the variation in gene expression comparable to that inferred from models of gene regulation based on the TFBS repertoire of core promoters. This observation underlines that TEs should not be ignored when attempting to delineate the regulatory programs orchestrating biological processes, in particular during embryogenesis.

We have also shown that *craTEs* identifies relevant cis-regulatory TE subfamilies with superior power compared to enrichment approaches based on differential expression. In addition, *craTEs* readily identifies TE-dependent cis-regulatory changes in experiments limited to a single replicate, whereas performing differential expression analysis requires at least an additional replicate in one of the conditions, i.e. three samples. This difference likely stems from how gene expression values are modeled in both methods. DE methods model the distribution of gene expression values across conditions independently for each gene, although current methods now leverage information borrowing techniques to share information across genes within samples [39]. In effect, DE methods perform one statistical test for each gene, resulting in tens of thousands of tests where the false discovery rate has to be controlled. Thus, any coordinated but mild difference in expression between co-regulated sets of genes is lost and cannot be used in the subsequent enrichment test. In contrast, *craTEs* leverages information across hundreds to thousands of genes to estimate the cis-regulatory activity of each TE subfamily in a single step.

Next, we empirically determined that the typical range until which cis-regulatory TEs regulate their target promoters is 200kb. To this end, we applied a cross-validation procedure to a large scale primed hESCs perturbation dataset to select the cis-regulatory distance that minimized the error between true and predicted gene expression values as estimated by *craTEs*. To our knowledge, this is the first attempt aiming at quantitatively estimating such distance by aggregating transcriptomic data derived from hundreds of experimental perturbations. Thus, TE subfamilies exert cis-regulatory influences up to distances compatible with those typically separating enhancers from their target promoters, consistent with the notion that many TEeRS are, in fact, bona fide enhancers.

We further characterized the landscape of TF overexpression-induced TE-dependent cis-regulatory changes. TFs poised towards the GRN of the naïve hESC state, namely TFAP2C, KLFs and NR5A1, collectively increased the cis-regulatory activity of LTR5-Hs, SVA and LTR7Y subfamilies, which function as KLF4-responsive enhancers in naïve hESCs. Whether these newly-identified inducers of evolutionarily young and cis-regulatory TE subfamilies mediate their effect via direct binding or secondary transcriptional changes will require further work. Still, these results underline the importance of LTR5-Hs, SVA and LTR7Y subfamilies in the GRN of naïve pluripotency. While the cis-regulatory activity of young TE subfamilies in pre-implantation embryogenesis is increasingly being recognized [63], the landscape of TE-dependent cis-regulation at later stages of human embryogenesis is still ill-defined. In the present study, we observed that overexpressing key regulators of gastrulation and germ-layer commitment – including GATA6 - in primed hESCs increased the cis-regulatory activity of LTR5-Hs and SVA subfamilies together with other primate-specific TE subfamilies. Importantly, we reported in a related manuscript that LTR5B, LTR5-Hs, LTR6A and LTR6B integrants are highly accessible in endodermal human fetal cells, more rarely in mesodermal fetal cells and generally inaccessible in ectodermal cells. In addition, we could confirm direct binding of GATA6 to LTR5B elements during hESC-derived endoderm differentiation [64]. Thus, the cis-regulatory role played by primate-restricted TEs during the pre-implantation stages of embryogenesis appears maintained, if not reactivated, during later stages embryogenesis.

Lastly, we leveraged epigenomics data to test whether changes in chromatin states and evidence for direct TF binding at TEs could single-out cis-regulatory integrants from non-cis-regulatory integrants within subfamilies. Surprisingly, we found that in the case of ZNF611 and KLF4 overexpression, TF binding was better able to enrich for cis-regulatory integrants than changes in histone marks. In the case of KLF4 overexpression, it is possible that the partial activation of compensatory TE-silencing mechanisms caused a divergence between the epigenomic and transcriptomic-derived cis-regulatory signals.

## Conclusion

That a simple mathematical model based on TE-promoter distances and the expression of protein-coding genes can infer cis-regulatory TE activities illustrates that as TEs spread, they rewire nearby protein coding genes into a web of regulatory dependencies which can be simultaneously fine-tuned by only a handful of transcriptional regulators. Furthermore, these recently emerged GRN components appear to regulate not only early embryogenesis, but also more advanced stages of development. For such vital and highly conserved events, the resulting speciation is only mechanistic owing to selective pressures. However, in situations allowing for more phenotypic diversification, for instance in the brain, the rapidly evolving TE-based cis-acting regulome likely contributes to the emergence of new traits.

## Methods

### Cell culture

#### Treatment protocol

Primed H1 were transduced with GFP or KLF4-containing lentiviral vectors and split after 48h then selected using blasticydin for the 3 following days. naïve WIBR3dPE hESC cells in KN/2iL media were transduced with GFP or ZNF611-containing lentiviral vectors, split after 96h, then selected for a couple of passages with blasticydin on irradiated Mouse Embryonic Blasticidin-resistant (MMMbz).

#### Growth protocol

Conventional (primed) human ESC lines were maintained in mTSER for H1 (Male) on Matrigel, for WIBR3 (Female) on irradiated inactivated mouse embryonic fibroblast (MEF) feeders in human ESC medium (hESM) and passaged with collagenase and dispase, followed by sequential sedimentation steps in hESM to remove single cells while naïve ES cells and primed H1 were passaged by Accutase in single cells. hES media composition: DMEM/F12 supplemented with 15% fetalbovine serum, 5% KnockOut Serum Replacement, 2 mM L-glutamine, 1% nonessential amino acids, 1% penicillin-streptomycin (Lonza), 0.1 mM β-mercaptoethanol and 4 ng/ml FGF2. naïve media composition: 500 mL of medium was generated by including: 240 mL DMEM/F12, 240 mL Neurobasal, 5 mL N2 supplement, 10 mL B27 supplement, 2 mM L-glutamine, 1% nonessential amino acids, 0.1 mM β-mercaptoethanol, 1% penicillin-streptomycin, 50 μg/ml BSA. In addition for KN/2i media: PD0325901 (1 μM), CHIR99021 (1 μM), 20 ng/ml hLIF and Doxycycline (2 μg/ml).

### ChIP-seq

Cells were cross-linked for 10 minutes at room temperature by the addition of one-tenth of the volume of 11% formaldehyde solution to the PBS followed by quenching with glycine. Cells were washed twice with PBS, then the supernatant was aspirated and the cell pellet was conserved in −80°C. Pellets were lysed, resuspended in 1mL of LB1 on ice for 10 min (50 mM HEPES-KOH pH 7.4, 140 mM NaCl, 1 mM EDTA, 0.5 mM EGTA, 10% Glycerol, 0.5% NP40, 0.25% Tx100, protease inhibitors), then after centrifugation resuspend in LB2 on ice for 10 min (10 mM Tris pH 8.0, 200 mM NaCl, 1 mM EDTA, 0.5 mM EGTA and protease inhibitors). After centrifugation, resuspend in LB3 (10 mM Tris pH 8.0, 200 mM NaCl, 1 mM EDTA, 0.5 mM EGTA, 0.1% NaDOC, 0.1% SDS and protease inhibitors) for histone marks and SDS shearing buffer (10 mM Tris pH8, EDTA 1mM, SDS 0.15% and protease inhibitors) for transcription factor and sonicated (Covaris settings: 5% duty, 200 cycle, 140 PIP, 20 min), yielding genomic DNA fragments with a bulk size of 100-300bp. Coating of the beads with the specific antibody and carried out during the day at 4°C, then chromatin was added overnight at 4°C for histone marks while antibody for transcription factor is incubated with chromatin first with 1% Triton and 150mM NaCl. Subsequently, washes were performed with 2x Low Salt Wash Buffer (10 mM Tris pH 8, 1 mM EDTA, 150mM NaCl, 0.15% SDS), 1x High Salt Wash Buffer (10 mM Tris pH 8, 1 mM EDTA, 500 mM NaCl, 0.15% SDS), 1x LiCl buffer (10 mM Tris pH 8, 1 mM EDTA, 0.5 mM EGTA, 250 mM LiCl, 1% NP40, 1% NaDOC) and 1 with TE buffer. Final DNA was purified with Qiagen Elute Column. Up to 10 nanograms of ChIPed DNA or input DNA (Input) were prepared for sequencing. Library was quality checked by DNA high sensitivity chip (Agilent). Quality controlled samples were then quantified by picogreen (Qubit 2.0 Fluorometer, Invitrogen). Cluster amplification and following sequencing steps strictly followed the Illumina standard protocol. Libraries were ligated with Illumina adaptors. Sequenced reads were demultiplexed to attribute each read to a DNA sample and then aligned to reference human genome hg19 with bowtie2. Peaks were called on mapped data using MACS2 [65]. Differential analysis between conditions has been performed with VOOM using unique reads (filter for MAPQ *<* 10), counted on the union of all peaks of a same experiment. Samples were normalized for sequencing depth using the counts on the union peaks as library size and using the TMM method [38] as it is implemented in the limma package of Bioconductor.

### RNA-seq analysis

#### Mapping

Reads were mapped to the human (hg19) genome using hisat2[**?**] with parameters `hisat2 -k 5 --seed 42 -p 7`.

#### Summarization

Counts on genes and TEs were generated using featureCounts [66]. To avoid read assignation ambiguity between genes and TEs, a gtf file containing both was provided to featureCounts. For repetitive sequences, an in-house curated version of the Repbase database was used (fragmented EREs belonging to the same subfamily were merged). Only uniquely mapped reads were used for counting on genes and TEs. Finally, features that did not have at least one sample with 20 reads were discarded from the analysis. Only features corresponding to protein-coding genes were kept.

#### Normalization

For input into *craTEs*, raw counts were transformed to transcripts per millions (TPM). A pseudocount equal to the fifth percentile of non-zero counts in the sample was added to each raw count before transformation to TPM and subsequent log_2_ transformation.

### Differential expression analysis-based cis-regulatory TE subfamily detection

DE analysis was performed using edgeR [38]. Starting from raw counts restricted to protein-coding genes, we performed library size normalization with the trimmed mean of M-values (TMM) normalization method [67]. We assumed that TMM-normalized counts follow a negative binomial distribution and estimated per-gene dispersions using the `estimateDisp` function from edgeR. We tested for differential expression using Fisher’s exact test as implemented in the function `exactTest` from edgeR. We either considered DE genes as those with Benjamini-Hochberg adjusted p values *<* 0.05 (stringent DE calling), or those with p values *<* 0.05 (lenient DE calling). Next, using the hypergeometric distribution, we computed for each TE subfamily the probability of finding more DE genes within cis-regulatory distance of its integrants than what was observed [37, 5]. We performed this last step separately for upregulated and downregulated genes. Finally, we gathered the results obtained for up/downregulated genes into a single table and accounted for multiple testing using the Benjamini-Hochberg procedure [34].

### cis-regulatory activity estimation for TE subfamilies (*craTEs*)

The *craTEs* model is adapted from the motif activity response analysis (MARA) model of gene regulation [28]. Let *E* be the matrix of gene expression, with *P* protein coding genes as rows, and *S* samples as columns. *E*_*ps*_ is the logged TPM expression value for gene *p* in sample *s*. Let *N* be the predictor/feature matrix with *P* protein coding genes as rows and *M* TE subfamilies as columns. *N*_*pm*_ is regulatory susceptibility [26] of protein-coding gene *p* to TE subfamily *m*, and in the absence of weighting procedure is computed as the number of times an integrant belonging to TE subfamily *m* is found in the vicinity of *p*. Let *A* be the matrix of cis-regulatory TE subfamily activities, with *M* TE subfamilies as rows and *S* samples as columns. *A*_*ms*_ is the cis-regulatory activity of TE subfamily *M* in sample *S. A*_*ms*_ can be seen as follows: if a TE integrant from subfamily *m* is inserted in the vicinity of gene *p*, the expression of gene *p* increases by the value *A*_*ms*_. Then, the expression *E*_*ps*_ of gene *p* in sample *s* is given by:

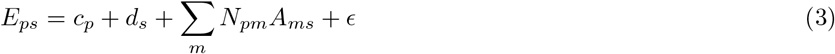

where *c*_*p*_ is a gene-specific constant representing basal transcription and *d*_*s*_ is a sample-specific constant that models sample-specific batch effects such as PCR amplification biases. The model across samples and genes can be written as

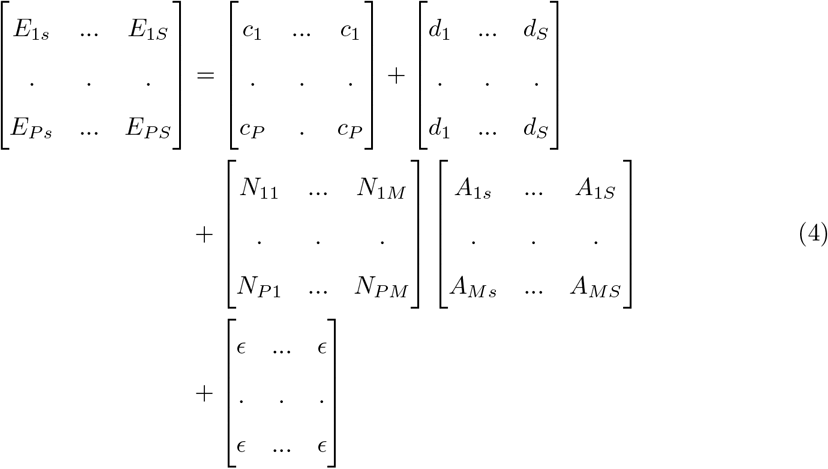

Column-centering *E* sets *d*_*s*_ to zero for each sample. Similarly, row-centering *E* sets *c*_*p*_ to zero for each gene. After row and column centering, the model becomes:

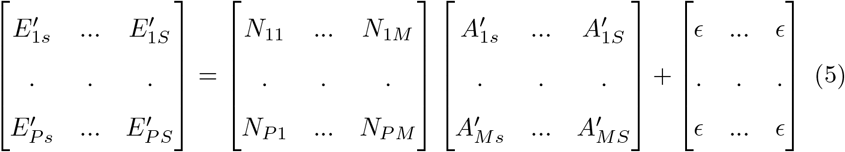

where 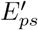 represents the deviation in expression from the average expression for gene *p* across all samples and 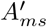 the deviation in cis-regulatory activity from the average cis-regulatory activity for gene *p* across all samples. The model is allowed to have a non-zero intercept, therefore the true model we fit is:

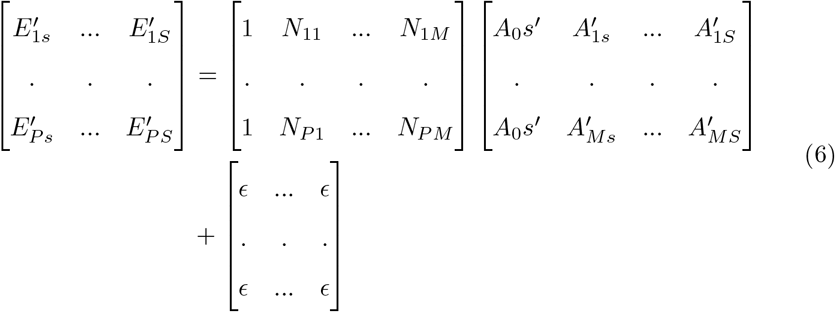

MARA [28] uses using ridge regression and picks the regularization parameter *λ* using 5-fold cross validation. *λ* controls for overfitting by imposing a so-called “budget” on TE activities. This method addresses the curse of dimensionality (too many predictors with respect to the number of observations) and stability issues arising when there is a high collinearity in the space of predictors. but the statistical significance of each predictor is more difficult to compute than in the standard linear regression setting. Additionally, in the MARA model, each activity deviates from a mean activity corresponding to an baseline regulatory state which can be hard to describe. Instead, we chose to consider samples in pairs. We contrasted samples from condition 2 (e.g. treatment samples) with samples from condition 1 (e.g. control samples). Under the normalized MARA-like model:

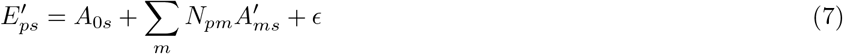

We are interested in contrasting two samples labeled sample 1 and sample 2.

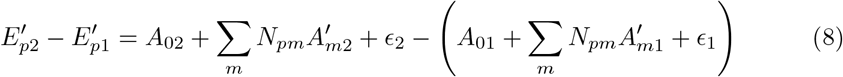

Therefore, we obtain:

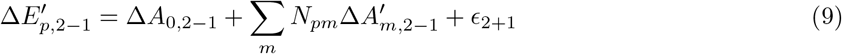

We used (9) as a model with identically and independently normal-distributed noise to estimate differences in activity between treatment and control samples. We then tested whether each estimated activity was t-distributed around 0. We controlled the false discovery rate using the Benjamini-Hochberg procedure. Paired replicates were treated by concatenating the vectors of differences in expression 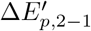 for each pair. The susceptibility matrix *N* was expanded row-wise accordingly.

### Computing the regulatory susceptibilities of each gene to TE subfamilies

The genomic locations of TEs was derived from Repeatmasker RELEASE 20170127, based on the hg19/GRCh37 assembly of the human reference genome. Repeat-Masker annotates TEs based on sequence similarity to a consensus sequence which tends to fragment partially degenerated integrants into multiple sequences. To avoid counting fragmented TEs several times, we merged TEs belonging to the same sub-family and the same strand separated by a genomic distance of less than 1 kb. The following procedure was applied to each protein-coding gene in order to delineate the set of corresponding putatively cis-regulatory integrants. The following steps were applied to each protein-coding gene (derived from ENSEMBL release 93 using Biomart) to designate the set of corresponding putatively cis-regulatory TEs. We defined gene promoter regions as clusters of transcription start sites (derived from ENSEMBL release 93 using Biomart) spaced by less than 1kb and extended by 500bp at their 5’ and 3’ ends. Next, we defined cis-regulatory windows as the union of promoter regions extended by 50kb at their 5’ and 3’ end. We identified all TEs present within cis-regulatory windows. We excluded TEs overlapping promoter regions as well as TEs overlapping exons. Finally, the remaining TEs were summed per subfamily to generate a vector representing the susceptibility of the gene to putatively cis-regulatory TEs.

### Building the susceptibility matrix N

The TE susceptibility matrix summarizes the potential regulatory activity of TE subfamilies on protein-coding genes. *N* was built by grouping integrants by sub-families and summing them for each gene. Therefore, *N*_*i,j*_ describes the number of integrants belonging to subfamily *j* in the cis-regulatory window of gene *i*.

### Weighting cis-regulatory TEs by their distance to gene promoters

To circumvent the need for a hard distance threshold, we weighted the regulatory potential of integrants by the distance separating them from gene promoters. Let *K* be the number of integrants of TE subfamily *m* present on the same chromosome as gene *p*. The regulatory potential of subfamily *m* on gene *p* is weighted by a Gaussian kernel: 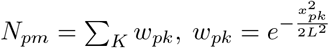 where:

1. *x*_*pk*_ is the distance in base pairs between the center coordinate of TE integrant *k* and the center coordinate of the closest promoter of gene *p*
2. *L* is the standard deviation (i.e. bandwidth) of the gaussian kernel, in base pairs

### Filtering *E* and *N*

Each experiment, defined as the set of treatment versus control expression vectors that will eventually form matrix *E*, was subjected to a separate filtering procedure. Genes with raw count values of less than 10 in all samples were removed from *E*. A per-column pseudo-count computed as the fifth percentile of all non-zero values in the column was added to each entry in *E. E* was transformed to transcript per millions (TPMs) and then log2-transformed. *E* was column-centered, and then row-centered. TE subfamilies with a sum of susceptibility scores Σ_*p*_ *N*_*pm*_ smaller than 150 were removed from *N*.

### Estimating the optimal TE-promoter regulatory distance

To estimate the optimal distance until which TE subfamilies regulate gene expression in cis, we built several weighted susceptibility matrices *N* by varying the values of *L* between 10^3^ and 10^1^0 base pairs and estimated the mean validation error using a 5-fold cross-validation on the gene space. The optimal value of *L* was chosen as the one that minimized the mean validation error. To ensure that that validation errors were comparable, we kept the sets of TE subfamilies and protein-coding genes fixed across all weighted matrices *N*. To this end, we filtered *E* and *N* according to the unweighted matrix *N* built with 100kB-wide cis-regulatory windows centered on gene promoters, as described above and in fig. 1. We then filtered each weighted susceptibility matrix *N* according to the rows (protein coding-genes) and columns (TE subfamilies) contained in the unweighted susceptibility matrix *N*.

### Splitting TE subfamilies between functional and non-functional fractions

Let *F* bet the set of genomic ranges considered as functional. Each TE integrant from subfamily *m* overlapping with at least one element in *F* was assigned to the so-called “functional” fraction of subfamily *m*: *m*_*functional*_. The matrix *N*_*functional*_ was built as described above for *N*, considering *m*_*functional*_ as a distinct subfamily. As splitting subfamilies into fractions may yield predictors, i.e. columns of *N*_*functional*_, with too few putatively regulated genes to reliably estimate TE sub-family cis-regulatory activities, we applied the following procedure:

- TE subfamilies (including their functional fractions) that were excluded by the filtering procedure applied on *N* described above were also excluded from *N*_*functional*_.

- If either the functional or the non-functional fraction of a TE subfamily showed Σ_*p*_ *N*_*pm*_ *<* 100, both fractions were removed and replaced with the corresponding column in *N*, i.e. the vector of regulatory susceptibility scores *N*_*pm*_ for the entire subfamily.

- We allowed some user-specified subfamilies to be “protected” from this filtering step. These subfamilies remained split between a functional and a non-functional fraction in *N*_*functional*_ irrespective of the sum of their regulatory susceptibility scores.

### Statistical methods

The statistical significance of TE subfamily activities is evaluated through null hypothesis testing, where the null hypothesis is *H*0 : the value of the associated linear regression coefficient *β* is zero. All p-values reported in the manuscript are adjusted for multiple testing using the Benjamini Hochberg procedure, except when specified in the methods or main text. We reject the *H*0 when the adj. p-value *≤* 0.05.

## Declarations

### Ethics approval and consent to participate

hESC usage has been approved by the Swiss Federal Office of Public Health, the Canton of Vaud Ethics Committee (Autorization Number R-FP-S-2-0009-0000) and registered in the European Human Pluripotent Stem Cell Registry (hPSCreg).

### Consent for publication

Not applicable.

### Availability of data and materials

The TE annotation database RepeatMasker library RELEASE 20170127 can be found on the RepeatMasker website accessible at URL http://repeatmasker.org/libraries/RepeatMaskerMetaData-20170127.tar.gz.

The following RNA-seq datasets: naïve hESCs + CRISPRi against SVA/LTR5-Hs, primed hESCs + GFP or KLF4, naïve hESCs + GFP or ZNF611; ATAC-seq dataset: naïve hESCs + CRISPRi against SVA/LTR5-Hs; ChIP-seq datasets: H3K9me3 in naïve hESCs + CRISPRi against SVA/LTR5-Hs, H3K9me3/H3K27ac in primed hESCs + GFP or KLF4, H3K9me3/H3K27ac in naïve hESCs + GFP or ZNF611 can be found on the Gene Expression Omnibus (GEO) under accession number GSE117395 at URL: https://www.ncbi.nlm.nih.gov/geo/query/acc.cgi?acc=GSE117395 [5].

The RNA-seq dataset of K562 + CRISPRi against LTR2B can be found on the Gene Expression Omnibus (GEO) under accession number GSE136763 at URL https://www.ncbi.nlm.nih.gov/geo/query/acc.cgi?acc=GSE136763 [32].

The RNA-seq dataset of transgene overexpression in hESCs can be found on the DNA Data Bank of Japan (DDBJ) Sequence Read Archive (DRA) under SRA submission number DRA006296 at URL https://ddbj.nig.ac.jp/resource/sra-submission/DRA006296 [40].

The following ChIP-seq datasets: ChIP-seq against KLF4 in primed hESCs, ChIP-seq against ZNF611 in naïve hESCs can be found on the Gene Expression Omnibus (GEO) under accession number GSE208403 at URL https://www.ncbi.nlm.nih.gov/geo/query/acc.cgi?&acc=GSE208403.

The regulatory susceptibility matrix N, TEs vs promoters, 100kB-wide windows can be found on ZENODO at URL https://doi.org/10.5281/zenodo.6707955 The regulatory susceptibility matrix N, TEs vs promoters, weighted with *L* = 1*e*5*kB* can be found on ZENODO at URL https://doi.org/10.5281/zenodo.6769794 The regulatory susceptibility matrices N with functional fractions, TEs vs promoters, weighted with *L* = 1*e*5*kB* can be found on ZENODO at URL https://doi.org/10.5281/zenodo.6769794

The regulatory susceptibility matrices N, TEs vs promoters, weighted with *L* in [1*e*3*kB*, 1*e*10*kB*] can be found on ZENODO at URL https://doi.org/10.5281/zenodo.6818283

The code used to process the data and generate the figures can be found and executed directly from the renkulab platform for reproducible data science at URL https://renkulab.io/gitlab/crates.

### Author’s contributions

CP designed the research plan, analyzed the data and wrote the manuscript with biological supervision by DT, JP and statistical supervision by RF. DT, JP and RF all made substantial contributions to the manuscript. JP generated the ChIP-seq data. DG, JD, SS and EP transformed the raw RNA-seq data into count tables, processed the ChIP-seq data, provided the code to merge fragmented EREs and provided the corresponding paragraphs in the Methods section.

## Acknowledgements

We thank Charlène Raclot and Sandra Offner for technical support regarding wet lab experiments; Alexandre Coudray, Eunji Shin, Paola Malsot and Felix Naef for scientific discussions; Nicolas Barrière, Cyril Matthey-Doret and the whole renku team for technical support regarding the renku platform and Séverine Reynard for administrative assistance.

## Additional files

### Additional file 1

`additional file 1.csv`, related to fig. 3 contains estimated TE subfamily activities for the RNA-seq dataset of transgene overexpression in hESCs [40] in .csv format, available at URL: https://doi.org/10.5281/zenodo.6826370. Title of data: Transposable element (TE) subfamily cis-regulatory activities estimated from 441 transgene overexpression experiments in human embryonic stem cells (hESCs).

Description of data: Activities were estimated using *craTEs* with the weighted susceptibility matrix N computed with *L* = 1*e*5*kB*. Rows are transposable element (TE) subfamilies, columns are as follows:

- Estimate: estimated cis-regulatory activity, correponds to a linear regression coefficient
- Std. Error: standard error of linear regression coefficient
- t value: t value corresponding to t-test with H0: Estimate = 0 and HA: Estimate ≠ 0
- *Pr*(*>* |*t*|) : probability (p-value) of observing a more extreme t value
- p_adj: p-values adjusted with the Benjamini Hochberg procedure
- TE: TE subfamily
- condition: concatenation of the name of the overexpressed transgene and the timepoint
- transgene: symbol for the overexpressed transgene
- timepoint: time under transgene induction via DOX treatment

### Additional file 2

`additional file 2.pdf`, related to fig. 3 contains the full heatmap of TE subfamily cis-regulatory activity statistical strengths (-log10 adj. p-value) estimated from dox-induced TF overexpression experiments in primed hESCs[40] as described in the legend of fig. 3B, with row and column dendrograms.

## Figures

**Figure S1.**
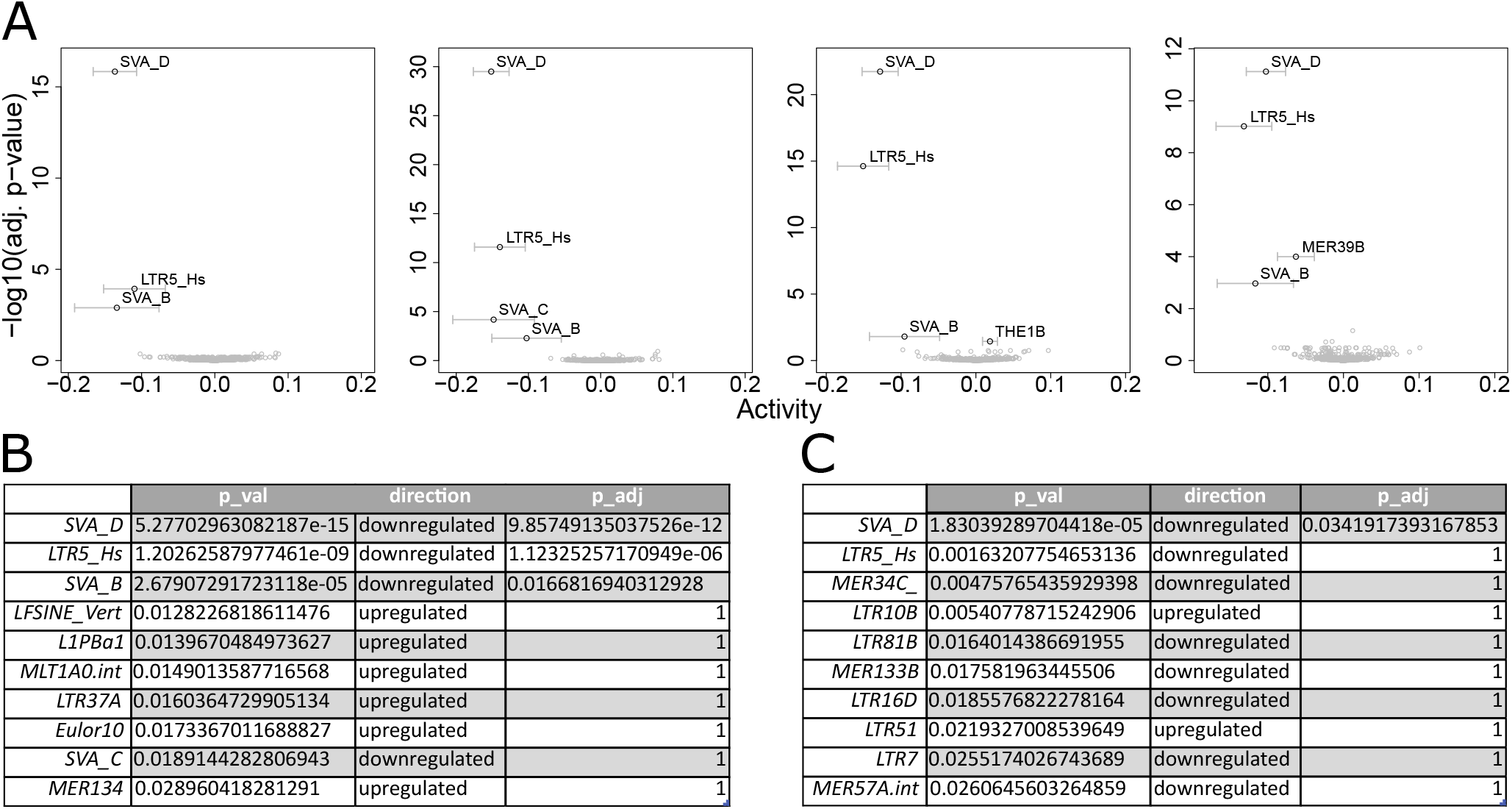
craTEs outperforms enrichment approaches based on differential expression analyses. **A** Case study for the estimation of the TE subfamily cis-regulatory activities in a 1 vs 1 sample setting (*n* = 1), using each of the paired replicates in the CRISPRi-mediated repression of LTR5-Hs and SVAs in naïve hESCs. **B** Top enriched TE subfamilies found in the proximity of differentially expressed (DE) genes. The sign of the log fold change (up/downregulation) is reported in the “direction” column. DE genes were called without adjusting for multiple testing. p-values for the enrichment test were computed according to the hypergeometric distribution, and adjusted with the Benjamini-Hochberg procedure[34]. **C** Top enriched TE subfamilies found in the proximity of DE genes, called with p-value adjustment using the Benjamini Hochberg procedure.

**Figure S2.**
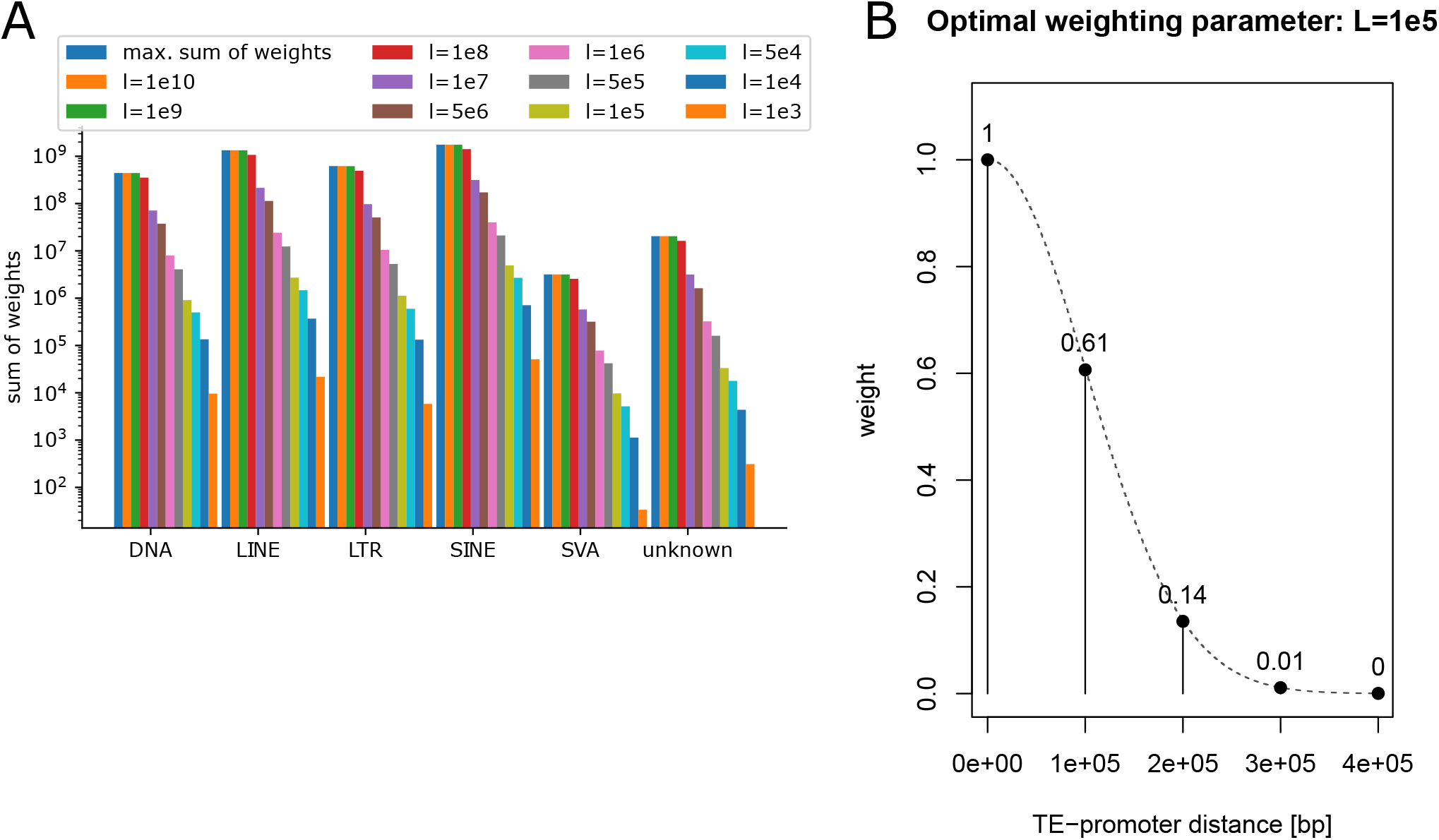
Related to Figure 2. **A** Sum of cis-regulatory weights as a function of the gaussian kernel width *L* across each main TE families. The maximum cis-regulatory weight represents the limit case whereby each TE contributes to the regulation of each gene located on the same chromosome. **B** Illustration of the gaussian kernel corresponding to the optimal choice of *L* = 100kb.

