## Supplementary figures and images for "Statistical learning quantifies transposable element-mediated cis-regulation"

### additional file 2

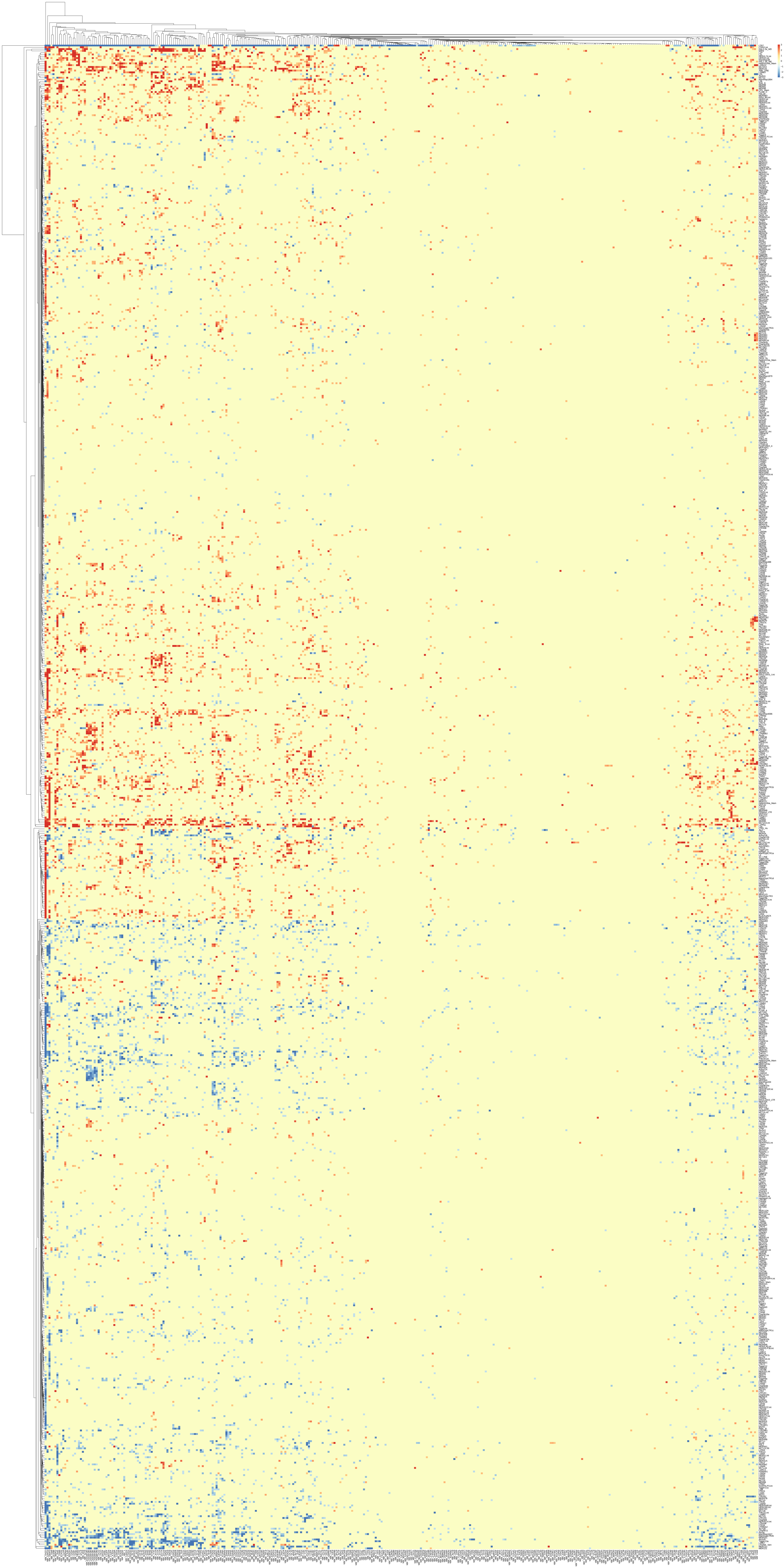
